# The Antarctic Seafloor Annotated Imagery Database

**DOI:** 10.1101/2023.02.16.528770

**Authors:** Jan Jansen, Victor Shelamoff, Charley Gros, Thomas Windsor, Nicole A. Hill, David K. Barnes, David A. Bowden, Julian Gutt, Narissa Bax, Rachel Downey, Marc Eléaume, Alexandra L. Post, Huw Griffiths, Katrin Linse, Dieter Piepenburg, Autun Purser, Craig R. Smith, Amanda F. Ziegler, Craig R. Johnson

**Affiliations:** Institute for Marine and Antarctic Studies, University of Tasmania, Hobart, AUS; British Antarctic Survey, UKRI, Cambridge, UK; National Institute of Water and Atmospheric Research, Wellington, NZ; Alfred Wegener Institute, Helmholtz Centre for Polar and Marine Research, Bremerhaven, GER; South Atlantic Environmental Research Institute, Stanley, FLK; Centre for Marine Socioecology, Institute for Marine and Antarctic Studies, University of Tasmania, Hobart, AUS; Fenner School of Environment and Society, Australia National University, Canberra, AUS; Muséum National d’Histoire Naturelle, UMR 7205-ISYEB, Centre National de la Recherche Scientifique-UPMC-EPHE, Paris, FRA; Geoscience Australia, Canberra, AUS; Department of Oceanography, University of Hawai’i at Mānoa, Honolulu, USA; Department of Arctic and Marine Biology, The Arctic University of Norway, Tromsø, NOR

**Author notes:** corresponding author: Jan Jansen.

## Abstract

Marine imagery is a comparatively cost-effective way to collect data on seafloor organisms, biodiversity and habitat morphology. However, annotating these images to extract detailed biological information is time-consuming and expensive, and reference libraries of consistently annotated seafloor images are rarely publicly available. Here, we present the Antarctic Seafloor Annotated Imagery Database (AS-AID), a result of a multinational collaboration to collate and annotate regional seafloor imagery datasets from 19 Antarctic research cruises between 1985 and 2019. AS-AID comprises of 3,599 georeferenced downward facing seafloor images that have been labelled with a total of 615,051 expert annotations. Annotations are based on the CATAMI (Collaborative and Automated Tools for Analysis of Marine Imagery) classification scheme and have been reviewed by experts. In addition, because the pixel location of each annotation within each image is available, annotations can be viewed easily and customised to suit individual research priorities.

This dataset can be used to investigate species distributions, community patterns, it provides a reference to assess change through time, and can be used to train algorithms to automatically detect and annotate marine fauna.

## Background & Summary

Marine imagery is a cost-effective and non-intrusive way to gather information about the seafloor. Extracting sound and detailed biological information from these images, however, is a key bottleneck because image annotation is time-consuming and often expensive. Perhaps because these data are difficult to produce, they have historically rarely been shared on public repositories. There has been a recent societal and scientific shift towards sharing data, greater transparency and reproducibility of data (FAIR data principles)^1,2^, and increasing capabilities of machine learning methods that require large numbers of annotated images has encouraged more scientists and research institutions to make datasets of expert annotated marine images publicly available (e.g., FathomNet^3^). Nonetheless, for most regions around the world, including the Southern Ocean, expert annotated seafloor images are rarely publicly available.

The sparsity of biological data impedes efforts to benchmark the status of marine ecosystems, and therefore also prevents monitoring of ecological change. This is particularly evident for the highly biodiverse Antarctic seafloor^4–6^, which still lacks a comprehensive analysis of the distribution of its biodiversity on a continental scale. While Southern Ocean ecosystems are predicted to change significantly as global temperatures change^7–11^, the information underlying this knowledge stems from either single regions^8,10,12^ or global models^11^, from single taxonomic groups^9^ or from datasets that compile opportunistically collected fauna^7^. A database of the distribution and abundances of a broad range of organisms covering the entire Antarctic continental shelf is invaluable to help identifying key conservation areas and managing these unique ecosystems.

In this article, we introduce a database of expert annotated seafloor imagery from the Antarctic continental shelf and slope, the Antarctic Seafloor Annotated Imagery Database (AS-AID, Fig. 1). AS-AID comprises of 3,599 annotated images and 615,051 annotations of a total of 180 faunal classifications. Images have been collected from scientific surveys between 1985 and 2019 for which data was either publicly available or could be sourced by directly contacting data curators. This represents the first international effort to collate images from different regions of the Southern Ocean at a pan-Antarctic scale into a single database and annotate these images using a single classification scheme.

**Figure 1:**
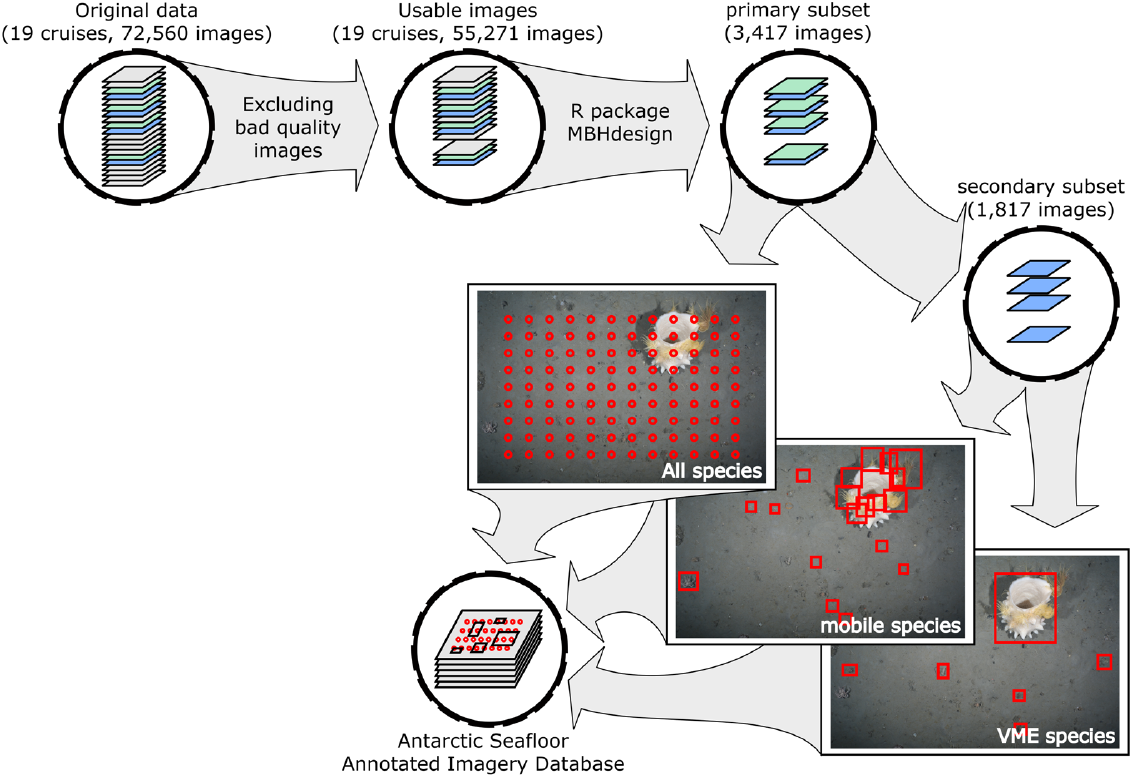
Simple overview of the steps involved in building AS-AID, including the methodology used to generate the database. Seabed photo sourced from pangea.de

Repeatability is key to making data useful beyond the scope of a single project, which is why annotations in AS-AID are based on an internationally recognised standard for labelling marine images (CATAMI^13^), and all annotations have exact reference points (x and y coordinates) to their respective images. Users can view AS-AID annotations on Squidle+ (www.squidle.org) and review all annotations for a single label and customise annotation labels to suit their own research objective. AS-AID can be used to train deep learning algorithms for automated classification of marine fauna. Combining AS-AID with environmental data and statistical models can allow insights into the species distributions and ecological processes that shape Antarctic seafloor ecosystems^10,14–16^.

## Methods

The Antarctic Seafloor Annotated Image Database (AS-AID) includes downward facing images of the seafloor at a depth range between 100 – 3,000 m from Antarctic scientific surveys for which we were able to obtain access to both images and detailed metadata records. We restricted the database to images deeper than 100 m because at these depths environmental conditions are more stable (e.g., relatively less impact on the fauna from iceberg scour) and more comparable between regions, which suits the continental scale biodiversity analysis that this database has primarily been developed for. AS-AID focusses on downward facing seafloor imagery rather than oblique facing imagery (where cameras can be mounted at different angles) because initial work indicated that downward facing images are more comparable between different surveys, with greater uniformity of illumination and resolution. Additionally, the area of seabed imaged with downward facing cameras can be more accurately estimated than from data collected with obliquely mounted cameras. In total, we identified 19 surveys from 2019 and prior that collected downward facing images of suitable quality with accessible metadata records also available (Fig. 1, 2 & Table 1). The only surveys that we identified as potentially suitable, but for which we could not access all relevant data were LMG1703 and ANA08D, which are therefore excluded from AS-AID at time of writing.

**Figure 2:**
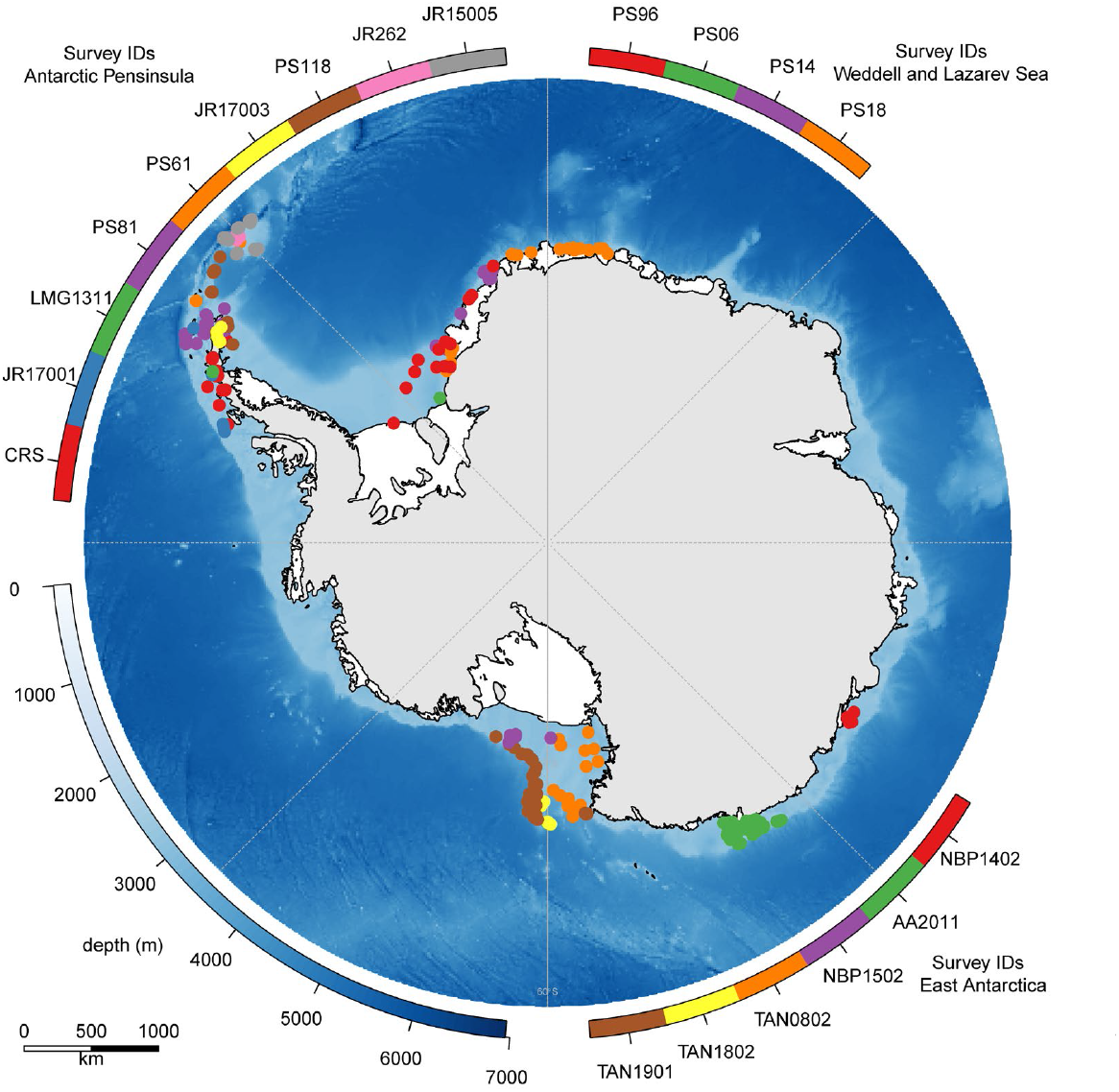
Overview of the circum-Antarctic distribution of sampling locations. Details about each survey can be found in tables 1-4.

**Figure 3:**
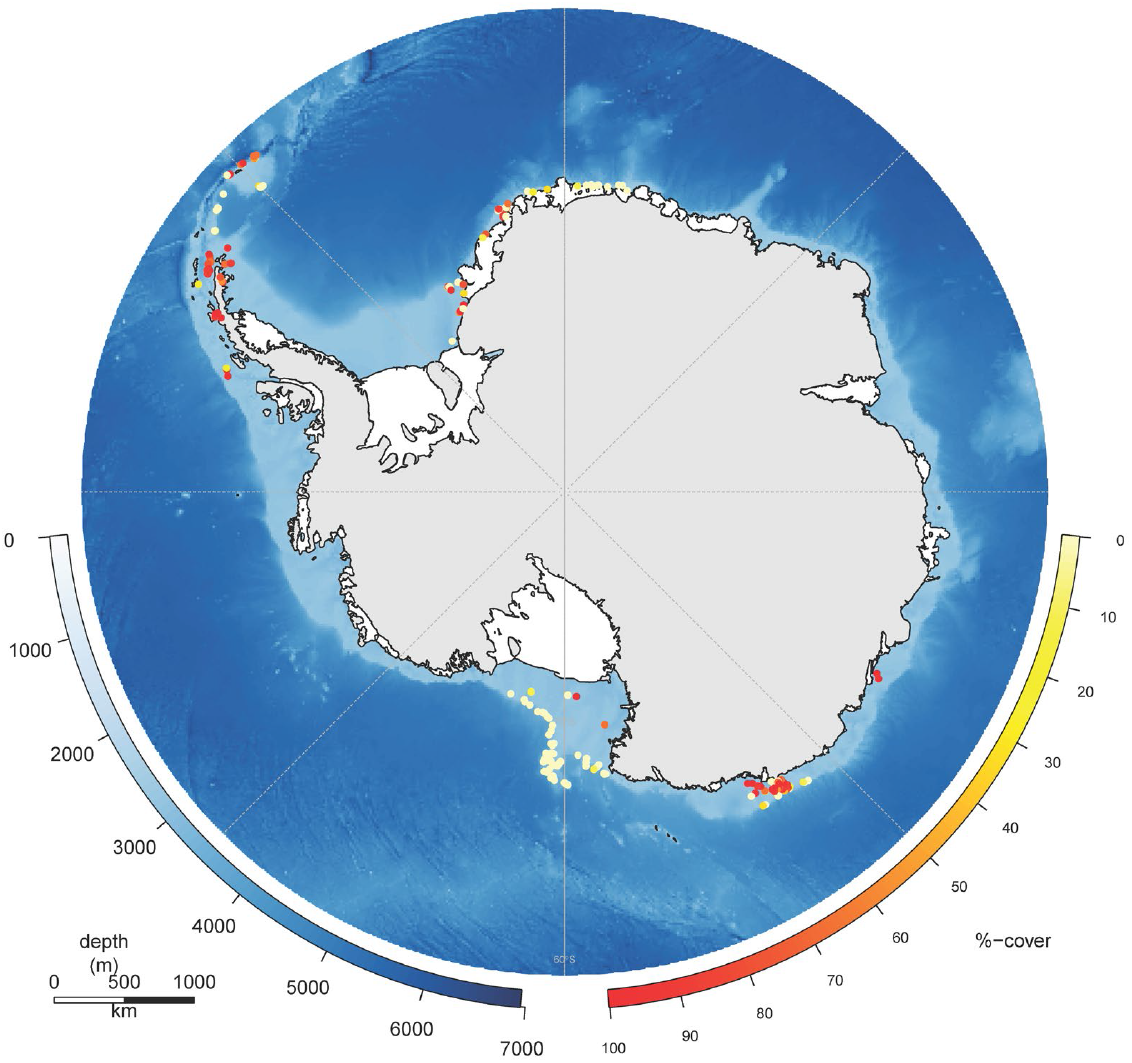
Percentage cover of fine substrate across all sampling locations

**Figure 4:**
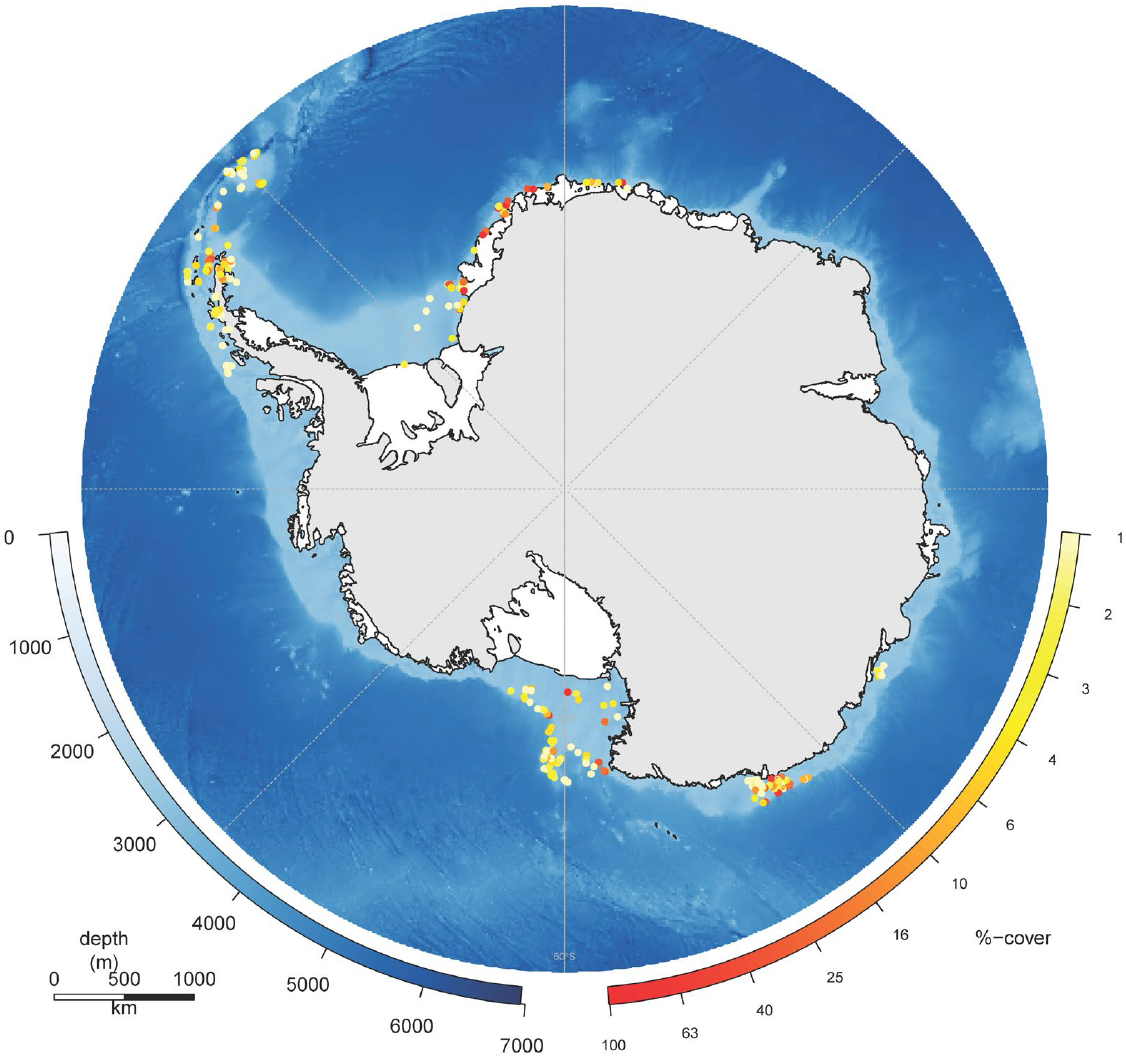
Combined percentage cover of all ascidians, bryozoans, cnidarians, hydrocorals and sponges at each sampling location.

**Table 1:**
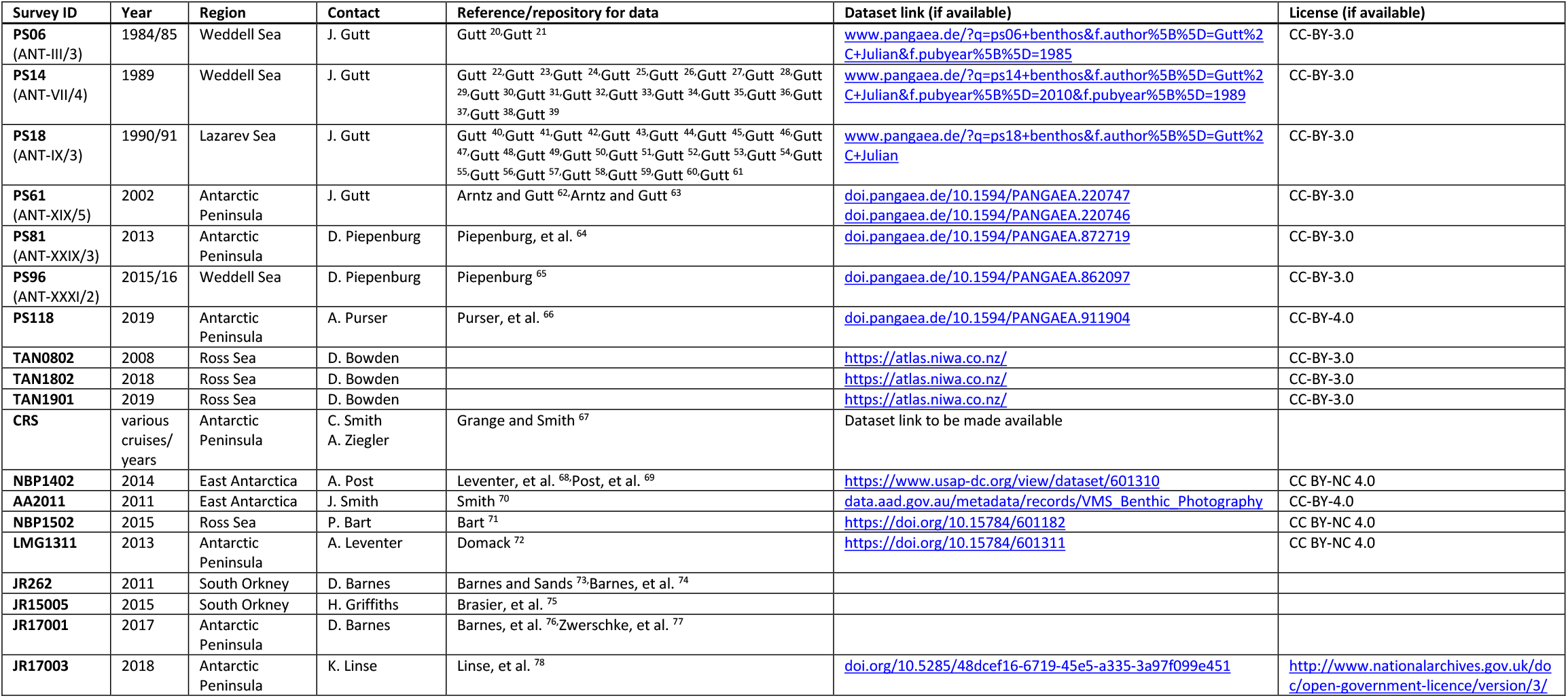
Survey details, including links and references to data repositories

### Sampling locations and image collection

AS-AID images were collected during 19 scientific cruises between 1985 and 2019 (Tables 1-4), using a range of different camera platforms (Table 4). Figure 2 shows an overview of sampling locations for all image transects included in AS-AID. Images and metadata were downloaded and stored on external hard-drives, and image filenames were standardised for processing.

**Table 2:**
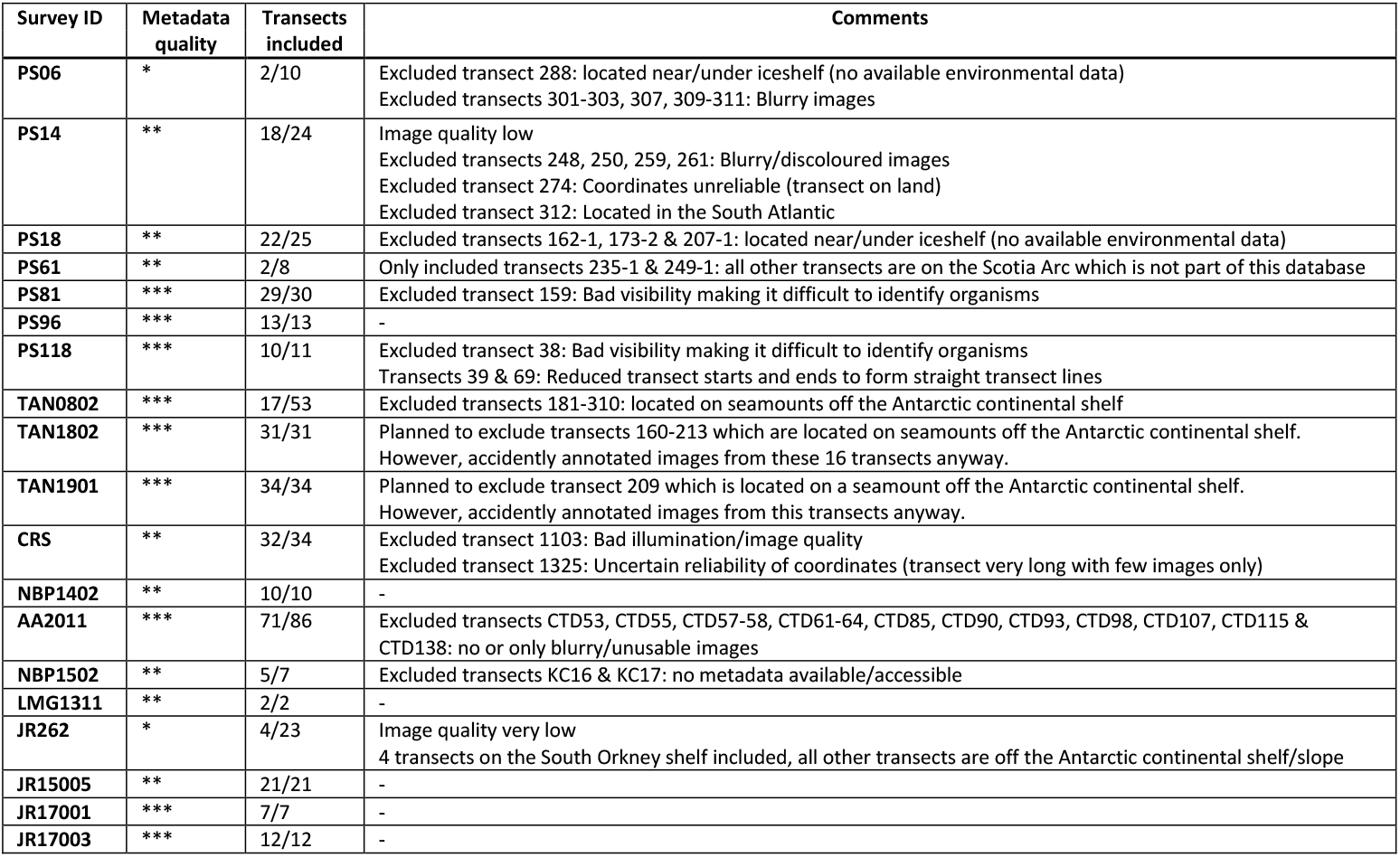
Data characteristics for each cruise and comments on which data were excluded. Metadata quality: * single lon/lat position for each transect; ** lon/lat available for start/end locations of each transect Locations of individual images interpolated from start/end locations; *** lon/lat available for each image

**Table 3:** Overview of the number of images per survey and how subsets were created. “Total” is the number of images downloaded from the repository; “Usable” is the number of images that show the seafloor, are in focus and are neither blurry nor too bright/dark; “Included” is the number of images added to our database, that is images that are from transects that we were interested in analysing; and “Subset 1” and “Subset 2” are the number of images that were selected for annotation for the point-scoring method and the exhaustive search respectively.

| Survey ID | Number of images |  |  |  |  | Image subsetting |
| --- | --- | --- | --- | --- | --- | --- |
|  | Total | Usable | Included | Subset 1 | Subset 2 |  |
| PS06 | 500 | 31 | 16 | 9 | 5 | Transect 289: selected 5 random images<br>Transect 292: selected all good quality images (4) |
| PS14 | 1523 | 1220 | 1098 | 119 | 65 | Spatially-balanced-random<br>However: Start/end coordinates not trustworthy (ship speed would be >1kt) for transects 245, 250, 256, 259, 260, 261, 275, 278-3, 280 & 293: Selected 5 random images out of the first 25 images from these transects, using starting GPS coordinates as image position. |
| PS18 | 1705 | 1654 | 1456 | 103 | 59 | Spatially-balanced-random<br>However: Start/end coordinates not trustworthy (ship speed would be >1kt) for transects 171-1, 173-1, 206-2, 211-2 & 222-1: Selected 5 random images out of the first 25 images from these transects, using starting GPS coordinates as image position. |
| PS61 | 139 | 129 | 129 | 12 | 7 | Spatially-balanced-random |
| PS81 | 14543 | 14516 | 14021 | 1041 | 528 | Spatially-balanced-random |
| PS96 | 2736 | 2727 | 2727 | 217 | 112 | Spatially-balanced-random |
| PS118 | 12629 | 6057 | 4691 | 260 | 133 | Spatially-balanced-random<br>Transect 11-2: GPS-coordinates wiggle a lot between images -> subset based on distance from start rather than distance along transect<br>Transect 39-1: removed the first 800 and the last 487 images to result in a straight lined transect<br>Transect 69-1: removed the first 29 and the last 50 images to result in a straight lined transect |
| TAN0802 | 13046 | 11403 | 3310 | 173 | 91 | Spatially-balanced-random |
| TAN1802 | 7470 | 5616 | 5616 | 336 | 175 | Spatially-balanced-random |
| TAN1901 | 8301 | 5878 | 5878 | 373 | 195 | Spatially-balanced-random |
| CRS | 2396 | 2032 | 1982 | 320 | 169 | Spatially-balanced-random |
| NBP1402 | 761 | 692 | 453 | 78 | 43 | Spatially-balanced-random |
| AA2011 | 1853 | 435 | 435 | 71 | 71 | Selected 1 random image per transect due to low number of usable images |
| NBP1502 | 2869 | 1719 | 1719 | 245 | 124 | Spatially-balanced-random |
| LMG1311 | 263 | 222 | 222 | 60 | 31 | Spatially-balanced-random |
| JR262 | 61 | 31 | 31 | 17 | 10 | Selected 5 random images per transect |
| JR15005 | 773 | 505 | 505 | 88 | 53 | Spatially-balanced-random |
| JR17001 | 142 | 120 | 120 | 14 | 8 | Spatially-balanced-random |
| JR17003 | 850 | 279 | 279 | 63 | 33 | Spatially-balanced-random |
| TOTAL | 72560 | 55271 | 42090 | 3599 | 1912 |  |

**Table 4:** Details on image collection, image characteristics and annotation

| Survey ID | Camera-platform | Depth-range (IBCSO V2)<br>for images |  | Distance from seafloor | Size<br>of imaged<br>seabed area | Laser-points | Image<br>crop | Colour-<br>corrected | Point-score<br>all species | Annotations<br>Exhaustive search |  |
| --- | --- | --- | --- | --- | --- | --- | --- | --- | --- | --- | --- |
|  |  | included | annotated |  |  |  |  |  |  | mobile species | VME species |
| PS06 | Photo sledge (FTS) | 349 – 1115 | same | Constant (weight) | Unknown | - | No | Yes | 972 | 241 | 10 |
| PS14 | Photo sledge (FTS) | 148 – 1347 | same | Constant (weight) | 0.56m<sup>2</sup> | - | No | Yes | 12852 | 1110 | 119 |
| PS18 | Photo sledge (FTS) | 144 – 956 | 176 – 956 | Constant (weight) | ~0.9m<sup>2</sup> (but cruise report says 1m<sup>2</sup>...) | - | No | Yes | 11124 | 1889 | 180 |
| PS61 | Photo sledge (FTS) | 330 – 348 | same | Constant (weight) | ~1m<sup>2</sup> | - | No | Yes | 1296 | 218 | 4 |
| PS81 | OFOS | 52 – 783 | same | Variable | Calculate | 50cm | 80% | Yes | 112428 | 83888 | 15249 |
| PS96 | OFOS | 198 – 731 | same | Variable | Provided + calculated | 50cm | 80% | No | 23436 | 7602 | 3852 |
| PS118 | OFOBS | 82 – 2169 | same | Variable | Calculate | 50cm | 80% | Yes | 28080 | 24900 | 6692 |
| TAN0802 | DTIS | 287 – 2258 | same | Variable | Calculate | 20cm | No | Yes | 18684 | 6237 | 872 |
| TAN1802 | DTIS | 664 – 1213 | same | Variable | Calculate | 20cm | No | Yes | 36288 | 12279 | 5964 |
| TAN1901 | DTIS | 447 – 1373 | same | Variable | Calculate | 20cm | No | Yes | 40284 | 16837 | 9523 |
| CRS | YOYO | 239 – 693 | same | Constant (weight, 2.5m) | ~3m<sup>2</sup> + calculated | 10cm | No | Yes | 34560 | 5850 | 573 |
| NBP1402 | YOYO | 281 – 899 | same | Constant (weight, 2.5m) | ~4.8m<sup>2</sup> + calculated | 10cm | No | Yes | 8424 | 2474 | 342 |
| AA2011 | CTD | 198 – 1453 | same | Variable | Calculate | 50cm | 90% | Yes | 7668 | 5322 | 543 |
| NBP1502 | YOYO | 175 – 597 | same | Calculate | Calculate | 10cm | No | Yes | 26460 | 6242 | 1281 |
| LMG1311 | YOYO | 349 – 654 | same | Calculate | Calculate | 10cm | No | Yes | 6480 | 1720 | 308 |
| JR262 | SUCS | 219 – 280 | same | Constant (frame) | 0.51m<sup>2</sup> | - | No | Yes | 1836 | 112 | 9 |
| JR15005 | SUCS | 473 – 1017 | same | Constant (frame) | 0.51m<sup>2</sup> | - | No | Yes | 9504 | 875 | 112 |
| JR17001 | SUCS | 67 – 507 | same | Constant (frame) | 0.51m<sup>2</sup> | - | No | Yes | 1512 | 74 | 21 |
| JR17003 | SUCS | 184 – 865 | same | Constant (frame) | 0.51m<sup>2</sup> | - | No | Yes | 6804 | 2702 | 133 |
| TOTAL |  | 67 – 2258 |  |  |  |  |  |  | 388692 | 180572 | 45787 |

### Image pre-processing

We inspected all 72,560 images visually and removed images/transects from AS-AID that were out of focus, too far or too close to the seafloor to allow identification of animals, had poor lighting, or had unreliable coordinates (Tables 2 & 3). We then corrected images for tone, colour and contrast using the automated batch function in Adobe Photoshop Version 21 (all surveys apart from PS96, see Table 4). Further, because the corners of images from surveys PS81, PS96, PS118 and AA2011 are out of focus we cropped these images to either 80% or 90% of their original size (Table 4).

### Image sub-setting

Annotating all images from all surveys is neither feasible nor desirable due to the large differences in sampling effort among surveys (i.e., the density of images along the transect lines is highly variable). We therefore created a subset of the full dataset by selecting a spatially balanced random subset of the images using their coordinates and the distance along the transect line with the R package MBHdesign version 2.2.2. Where only one coordinate for the transect was available, or where recorded end-coordinates of the transects did not match with expected values given the duration of the deployment and the normal ship-speed during deployment (max 1 kn), we randomly selected 5 of the first 25 images in those transects and assigned these images to the starting position of the transect (Table 2).

The primary image subset we created contains approximately 1 image for every 100 m of survey transect (3,599 images, Table 3). Further, we created a smaller secondary subset of images for a more detailed exhaustive search of each image, containing approximately 1 image for every 200 m of survey transect (1912 images, Table 3).

### Image annotation

We used both a point-grid and an exhaustive search for annotating the images. Point-grid annotation allows estimating percentage cover of all substrates and organisms present in the image, but the number of points that make up the grid influence how well these percentages are estimated, and how well rare and/or small organisms are detected^17^. In contrast, labelling organisms individually in an exhaustive search reduces detectability issues for rare species. However, individual labels present a challenge for many colonial organisms because individual colony sizes can vary greatly and it is frequently ambiguous how many colonies there are. We chose to first label all images in the primary subset using a point-grid, and then label the secondary subset using an exhaustive search. In the exhaustive search we focussed only on two groups of organisms: Mobile species, and small or rare species comprising Vulnerable Marine Ecosystem taxa (VME taxa as defined by the Commission for the Conservation of Antarctic Marine Living Resources, CCAMLR) that are not colonial.

For point-grid annotation we overlaid all images in the primary subset with a regular grid of 108 points (9×12) to closely match the 3×4 height to width ratio of the images. We chose 108 points as a good compromise between sampling effort and time constraints. For each point we only identified what substrate or organisms could be found within the single pixel in the centre of this point. While this required us to regularly zoom in and out of the image to determine the exact position of the point, it reduced potential bias towards labelling a point as an organism instead of substrate. We used CoralNet (coralnet.ucsd.edu/), which is a web-based toolkit with inbuilt deep-learning capabilities^18^. We manually annotated all images and used the inbuilt deep-learning algorithm as assistance to more quickly select the correct label.

For individually labelling organisms in the secondary subset via an exhaustive search, we used the online toolkit BIIGLE 2.0 (www.biigle.de/) and their “lawnmower mode” to systematically search through an image^19^. We identified each organism from the respective morphospecies list (Supplementary Classification Catalog) and drew a close-fitting circle around the organism if they were mainly (>50% of their body) within the image.

For all images in the primary subset, we used BIIGLE to annotate visible laser-points that allow calculating the imaged seabed area.

### Image classification scheme

We classified all organisms based on CATAMI^13^ (Collaborative and Annotation Tools for Analysis of Marine Imagery), and added labels to the classification tree when we could confidently distinguish organisms further into separate morphospecies.

CATAMI is a hierarchical classification scheme that uses taxonomy at the higher levels of the classification tree and morphology at the lower levels of the classification tree. This approach accounts for the fact that it is often difficult to identify seafloor fauna to a fine taxonomic resolution without physical specimens and that the morphology of seafloor fauna is often reflective of function. Where further discrimination is known or needed these can be added at the end of the respective CATAMI branch.

Our classification catalogue for both the point-scoring and the exhaustive search can be found in the supplementary information.

### Quality control & expert review

All annotations went through rigorous quality control comprising three stages: Labelling review, internal QA/QC, and expert review.

#### Labelling review concerns the exhaustive search only

After all images of a survey had been fully annotated in BIIGLE, a second scorer systematically searched through all images again to label any organisms the first scorer had missed. The use of two scorers to search though each image gives us high confidence that few individuals have been missed.

#### Internal QAQC

We conducted a two-stage internal QAQC both individually for each survey during annotation and also for all surveys combined. In the internal QAQC we reviewed each label, corrected any misclassification and discussed with the team if necessary. For reviewing labels from the grid annotation, we cropped 50×50 pixel thumbnails around each grid-point and sorted the thumbnails by their label in a morphospecies library. For reviewing the individual annotations from the exhaustive search using BIIGLE we used BIIGLE’s inbuilt “Largo” review function. After this first review, we then created a second library containing only thumbnails of morphospecies whose labels had changed during the review. This second review stage was to ensure all changes made to the original scoring were correct.

#### Expert review

In the third stage of our review process, we consulted with taxonomic experts to ensure the morphospecies grouping and labelling was consistent, and to help identify some of the unknown organisms encountered.

## Data Records

### Link 1

To be included here is a link to a persistent and publicly accessible repository that contains all images, metadata and annotations of AS-AID.

### Link 2

To be included here is a link to the data on SQUIDLE+. SQUIDLE+ has a user friendly interface that can be used to interrogate the annotated images in more detail.

## Technical Validation

Annotations in AS-AID have gone through a rigorous review process involving experienced scientists, trained students and taxonomists. During the annotation process, the annotation team (consisting of two scientists, one postgraduate student and one undergraduate student) met regularly (daily at the beginning, weekly at a later stage and every time a survey was fully annotated) to discuss uncertain annotations, ensure consistent annotation between the scorers and create new labels where necessary. All annotations were checked for correct labelling once a survey was fully annotated and the entire database was checked again after all surveys had been fully annotated. Further, expert taxonomists were consulted in particular for key taxa such as sponges, cnidarians and bryozoans. The largest uncertainties in this database were distinguishing whether an object was an organism or substrate, and whether an organism was dead or alive, and we created a number of categories to label these cases accordingly, such as e.g. “Unknown Biological Matrix” or “Unidentifiable”.

In AS-AID, we have opted to use experts to consecutively perform tasks on the same set of images. In particular, this meant one expert labelled the images while a second expert checked the annotations in the review process and flagged any misclassifications for discussion. While this process didn’t allow label accuracy to be quantified, it did allow organisms to be labelled more consistently throughout the entire database.

Researchers are invited to assess the validity of AS-AID for their own research purposes. Every annotation in AS-AID is linked to an image with a unique filename and contains the x- and y-coordinates where the organisms is found on that image as well as other metadata such as imaged area and imaging gear. Therefore, every annotation is fully traceable, allowing researchers to customise and assess the validity of AS-AID for their needs.

## Acknowledgements

We thank the crews and onboard scientific parties of the expeditions which collected the data within this study for providing the images and metadata for analysis in this study, either directly or via online raw data archives. We thank Amy Leventer for helping with data acquisition and Fanny Hermand for her help during the early stages of this project. This research has been funded as part of ARC Discovery Project 190101858. C.G is supported by a CSIRO-UTAS Quantitative Marine Science Scholarship. NZ involvement is supported by Ministry of Business Innovation and Employment programme C01X1710, *Ross Sea Research and Monitoring Programme; is the world’s largest MPA effective?*

## Author contributions

JJ: Conceptualization, Methodology, Software, Validation, Investigation, Data Curation, Writing - Original Draft, Writing – Review & Editing, Visualization, Supervision, Project Administration, Funding acquisition

VS: Methodology, Validation, Investigation, Data Curation, Writing – Review & Editing

CG: Methodology, Validation, Investigation, Data Curation, Writing – Review & Editing

TW: Methodology, Validation, Investigation, Writing – Review & Editing

NH: Conceptualization, Methodology, Writing – Review & Editing, Supervision, Project Administration, Funding acquisition

DKB: Investigation, Resources, Writing – Review & Editing

DAB: Investigation, Resources, Writing – Review & Editing

JG: Investigation, Resources, Writing – Review & Editing

NB: Investigation, Writing – Review & Editing

RD: Investigation, Writing – Review & Editing

ME: Investigation, Writing – Review & Editing

ALP: Investigation, Writing – Review & Editing

HG: Resources, Writing – Review & Editing

KL: Resources, Writing – Review & Editing

AP: Resources, Writing – Review & Editing

DP: Investigation, Resources, Writing – Review & Editing

CS: Resources, Writing – Review & Editing

AZ: Resources, Writing – Review & Editing

CJ: Conceptualization, Methodology, Writing – Review & Editing, Supervision, Project Administration, Funding acquisition

## Competing interests

The authors declare no competing financial interests.

## Supplementary information

The size of the classification catalog files is larger than the maximum allowed size for files at BioRxiv. The classification catalog files are available upon request from the corresponding author and will be made publicly available with the peer-reviewed version of this paper.

## Notes

### Competing Interest Statement

The authors have declared no competing interest.

